# Altered neural oscillations and behavior in a genetic mouse model of NMDA receptor hypofunction

**DOI:** 10.1101/2020.10.28.359547

**Authors:** David D. Aguilar, Leana K. Radzik, Felipe L. Schiffino, Oluwarotimi Folorunso, Mark R. Zielinski, Joseph T. Coyle, Darrick T. Balu, James M. McNally

## Abstract

**Introduction:** Abnormalities in electroencephalographic (EEG) biomarkers occur in patients with schizophrenia and those clinically at high risk for transition to psychosis and are associated with cognitive impairment. While the pathophysiology of schizophrenia remains poorly understood, converging evidence suggests *N*-methyl-D-aspartate receptor (NMDAR) hypofunction plays a central role and likely contributes to biomarker impairments. Thus, the characterization of such biomarkers is of significant interest for both the early diagnosis of schizophrenia and the development of novel treatments.

**Methods:** We utilized an established model of chronic NMDAR hypofunction, serine racemase knockout (SRKO) mice. *In vivo* EEG recording and behavioral analyses were performed on adult male and female SRKO mice and wild-type littermates to determine the impact of chronic NMDAR hypofunction on a battery of translationally-relevant electrophysiological biomarkers.

**Results:** SRKO mice displayed impairments in investigation-elicited gamma power that corresponded with reduced short-term social recognition. This impairment was associated with enhanced background (pre-investigation) broadband gamma activity that only appeared during social task performance. Additionally, SRKO mice exhibited sensory gating impairments, in both gamma power and event-related potential amplitude. However, other biomarkers such as the auditory steady-state response, sleep spindles, and state-specific power spectral density were generally neurotypical.

**Conclusions:** SRKO mice provide a useful model to understand how chronic NMDAR hypofunction contributes to deficits in a subset of translationally-relevant EEG biomarkers that are altered in schizophrenia. Importantly, our gamma band findings support the hypothesis that an aberrant signal-to-noise ratio impairing cognition occurs with NMDAR hypofunction, which may be tied to impaired task-dependent alteration in functional connectivity.

## INTRODUCTION

Abnormalities in neural network activity are common across a range of psychiatric disorders and may provide a diagnostic means for early diagnosis [1, 2]. A number of electroencephalographic (EEG) biomarkers are associated with impaired cognitive flexibility, plasticity and memory, executive functioning, and social behavior [3–6]. Of particular interest are disturbances in sensory gating and entrainment to 40Hz auditory stimuli (auditory steady state response; ASSR), which are impaired in patients with schizophrenia and individuals with clinical high risk to transition to psychosis [7]. Additionally, neural activity abnormalities have been reported in the gamma frequency range (30-80 Hz), either at rest or during cognitive and sensory related task performance [1].

Converging evidence suggests chronic NMDA receptor (NMDAR) hypofunction is central to the pathophysiology of schizophrenia and related psychiatric disorders [8, 9]. NMDAR antagonists can transiently recapitulate positive, negative, and cognitive symptoms of schizophrenia in healthy subjects and animal models [10, 11], including deficits in EEG biomarkers like ASSR [12]. These drugs also induce oxidative damage within cortical circuitry [13], potentially disrupting excitatory / inhibitory **(E/I)** balance and causing downstream abnormalities in gamma band oscillations and cognition. Indeed, patients with schizophrenia have well documented impairments in fast-spiking GABAergic interneurons, which may arise from these neurons’ increased susceptibility to oxidative damage [14–17]. Antipsychotic treatment can counteract NMDA antagonist-induced changes in gamma band oscillations, though their therapeutic mechanism is somewhat unclear [18]. Understanding the underlying mechanisms of EEG biomarkers could aid development of new targeted therapies that aim to alleviate cognitive deficits in patients with psychosis [19].

Recent genome-wide association studies have identified a number of genetic risk factors for schizophrenia associated with glutamatergic signaling, including the *SRR* gene which encodes serine racemase [20, 21]. Here we have examined the relationship between EEG biomarker activity and chronic NMDAR hypofunction by utilizing the serine racemase knockout (SRKO) mouse model. These mice lack expression of the enzyme responsible for synthesis of D-serine, a co-agonist at the NMDAR, and consequently exhibit chronic NMDAR hypofunction [22]. This well-established model exhibits a wide range of schizophrenia-like phenotypes [22–26] and exhibit enhanced oxidative damage and decreased parvalbumin immunoreactivity [27]. We hypothesized that SRKO mice will demonstrate impairments in translationally-relevant EEG biomarkers that are consistent with deficits associated with schizophrenia. The presence of these deficits may help us understand their underlying mechanisms and relationships to long term NMDAR dysfunction.

## MATERIALS AND METHODS

### Animals

SRKO mice were originally generated as described [22]. Adult male and female SRKO (-/-) mice and their wild type (WT) littermates were bred in-house from heterozygote SR (+/-) breeding pairs. These mice were maintained on a C57/BL6 background and were used for all experiments. Animals were given access to food and water *ad libitum* and maintained on a 12hr light / dark cycle (lights-on 7am). All procedures were performed in accordance with the National Institutes of Health guidelines and in compliance with the animal protocols approved by the VA Boston Healthcare System Institutional Animal Care and Use Committee.

### Stereotaxic Surgery

Adult (postnatal day 70+) mice were deeply anesthetized with isoflurane (5% induction, 1-2% maintenance) and body temperature was maintained with a chemical heating pad throughout the surgery. Epidural EEG screw electrodes (0.10”, Cat No. 8403, Pinnacle Technology Inc., Lawrence, Kansas, USA) were implanted in the skull above the frontal cortex (from bregma: A/P +1.9mm, M/L −1.0mm) and parietal cortex (A/P −1.0mm, M/L +1.0mm) with a reference and ground screw implanted above the cerebellum (from lambda, A/P −1.5mm, M/L ±1.3mm). Animals were given at least 7 days to recover from surgery before any experiments began. EEG/EMG signals were acquired via 3 channel amplifier (Pinnacle Technology), sampled at 2kHz, and low pass filtered at 200 Hz.

### Social Task-Elicited Gamma

We utilized a 3 chamber (Maze Engineers, Boston, MA, USA) task protocol modified from DeVito, Balu [25] described in **Figure 1 A**. For this experiment, video tracking software (Ethovision XT, Noldus) was used in combination with EEG (WinEDR, University of Strathclyde, Glasgow, Scotland) to record neural activity in freely behaving mice during investigations of the arena, an object, or another mouse (previously described in McNally, Aguilar [28], see **Supplemental Material A1**). EEG epochs were extracted around the investigation time, and gamma power (25-58Hz) was examined immediately after investigation onset vs a 4s “baseline” (0-4s pre-stimulus). This frequency range was selected because our preliminary data and previous studies [28] identified investigation-induced increases in 25-58Hz power, and this frequency range may signify top-down predictive coding in spatial working memory tasks [29]. Grand averages were taken across all epochs from all animals for each respective genotype after normalization.

**Figure 1.**
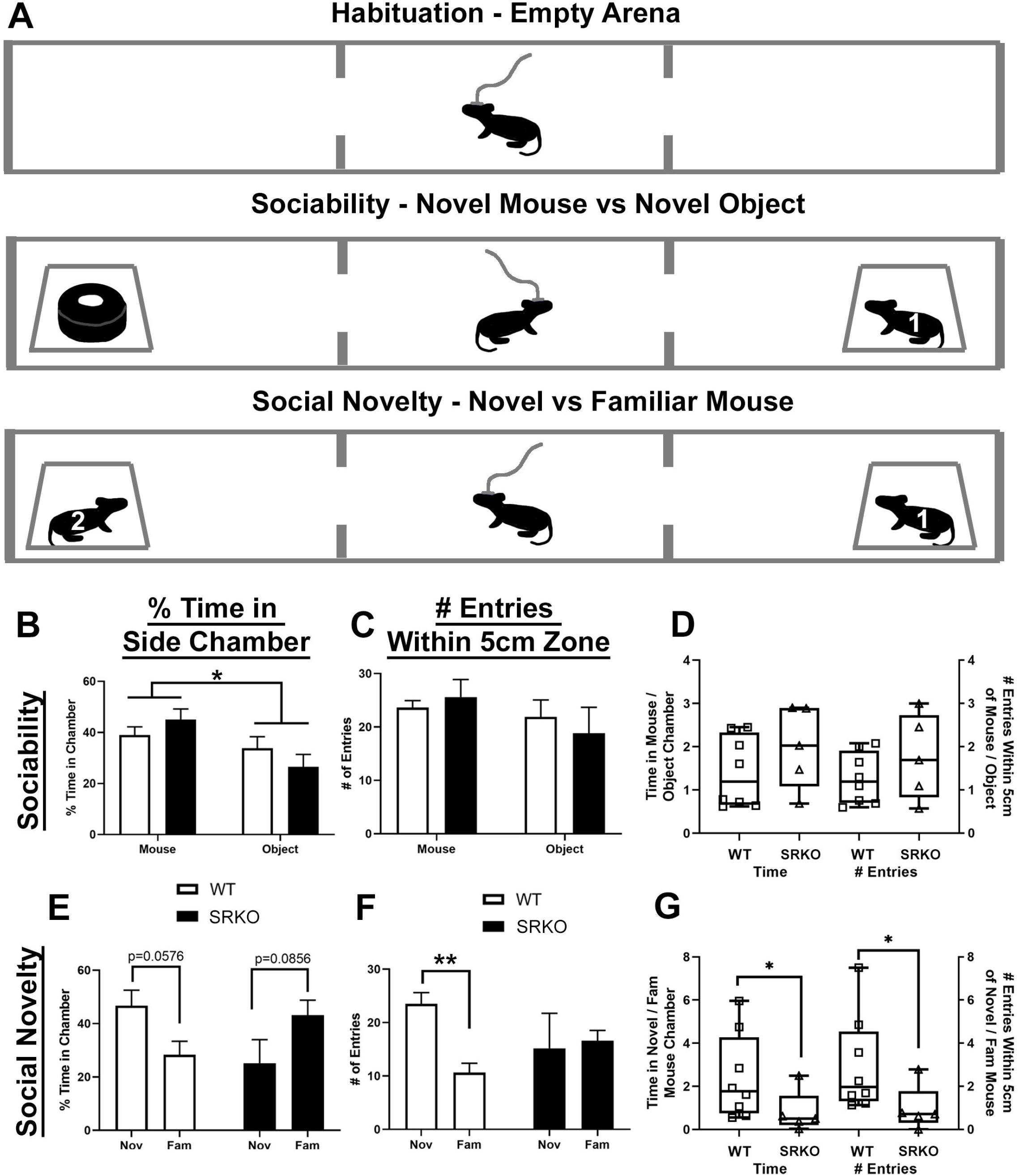
Social Novelty Recognition is Impaired in SRKO mice. Tethered, freely moving SRKO mice (n=5) and WT littermates (n=8) were placed in a three-chamber arena under low light conditions (14 lux) to measure sociability and social recognition following a 15 minute tethered habituation period in their home cage. A five minute habituation in the center chamber preceded each of the three consecutive stages shown in the diagram (**A**). During the empty arena habituation, animals had 5 minutes to explore all three empty chambers. During the 10 minute sociability stage, one side chamber contains an unfamiliar, older sex-matched mouse (stranger “1”) in a cage while the opposite chamber contains a similar looking novel object (e.g. a black roll of tape) in a cage. The location of the object and animal were counterbalanced between test animals. During the 10 minute social novelty stage the novel object was replaced with an unfamiliar sex-matched mouse (stranger “2”, age-matched to stranger 1) and the test animal could investigate the novel (“2”) and now familiar (“1”) mice. Behavioral analysis measured the proportion of time spent in each chamber and the number of entries within 5cm of each cage. During the sociability stage, both WT (white bars) and SRKO mice (black bars) spent a larger percent of time in the chamber containing the novel mouse than the chamber with the novel object (**B**) suggesting the sociability of SRKO mice is unchanged. There were no between-group differences during the sociability stage for either time spent exploring either chamber (**B**), number of entries within a 5cm zone surrounding the novel mouse / object (**C**) or in the novel mouse / object ratio of these measurements (**D**), indicating WT and SRKO mice behaved similarly during this phase of the task. During the social recognition stage only WT mice spent more time investigating the novel mouse than the familiar mouse, measured by the percent time in each chamber (**E**, strong trend) and the number of entries within a 5cm zone (**F**). There were significant between-group differences in the novel / familiar mouse ratio for both of these measurements (**G**), suggesting that WT animals spent a greater proportion of time investigating novel animals than familiar animals when compared to their SRKO littermates. This suggests SRKO mice demonstrate decreased social novelty recognition or impaired social novelty-related exploration. Stars represent a significant main effect (**B**) or a significant Holm-Sidak post hoc test following a significant interaction (**F**) in a two-way ANOVA. Stars in **G** represent a significant Mann-Whitney U test. In all figures, bar graphs represent mean values ± standard error of the mean (see **Supplemental Materials B1** for these values) while boxplots represent the median and 25^th^-75^th^ percentiles with whiskers that represent minimum and maximum values. Individual values are represented in **D** and **G** by hollow squares and triangles. For all experiments the number of stars represents the level of significance (* p<0.05, ** p<0.01, *** p<0.001, **** p<0.0001).

### Auditory Stimulation

Recordings occurred in each animal’s home cage within a sound-attenuated recording chamber (background noise ~55dB). Stimuli were generated by a BK Precision 4052 waveform generator (Yorba Linda, California, USA) using Spike2 software (Cambridge Electronic Design, Cambridge, UK) or WinWCP (University of Strathclyde, Glasgow, Scotland) and were delivered through speakers adjacent to the home cage. Data was collected with a Micro 1401 mkII interface module (CED, Cambridge, UK) and Pinnacle’s 3 channel amplifier.

### Sensory gating

Following a 10-minute tethered habituation period, auditory stimuli were presented as pairs of 80dB 5kHz tones of 50ms duration (n=100 trials) with an inter-trial interval (**ITI**) of 6s and a 500ms inter-stimulus interval (**ISI**). The average event-related potential (**ERP**) for the first (**S1**) and second (**S2**) stimulus were analyzed as described in Featherstone, Shin [30]. Briefly, a waveform average of 100 paired-tone presentations was created from raw EEG records. For each ERP, we measured the maximum positive deflection around 20ms (P20, 15-30ms after the tone) and the maximum negative deflection around 40ms (N40, 25-55ms) following a 100ms pre-stimulus baseline correction. See **Supplemental Material A2-A3** for more detail.

### ASSR

As the ASSR task yielded negative results, these methods and results appear in the **Supplemental Material** (**A4 and Table S1**).

### Data and Statistical Analyses

Data analysis was performed using custom scripts written for Matlab 2016a (Natick, MA, USA). Power spectral density (**PSD**) analysis was performed using the multi-taper method (social task-elicited gamma, resting state gamma, [31], Chronux Toolbox, chronux.org) or complex Morlet wavelet analysis (sensory gating, ASSR), as described previously [32]. See **Supplemental Material A2** for more detail on time-frequency spectral analysis. Statistical significance was set at p<0.05. Statistical analysis generally entailed a two-way ANOVA or two-way repeated-measures ANOVA with any significant interactions followed up by a Holm-Sidak multiple comparisons test. For comparisons between two groups, unpaired two-tailed Welch’s t tests were used. If the data failed a Kolmogorov-Smirnov test of normality, then a nonparametric analysis was run instead (Mann-Whitney). Repeated measures correlations were calculated in R using the rmcorr package [33]. Bar graphs represent mean values ± standard error of the mean (reported in **Supplemental Material B1**) while boxplots represent the 25^th^-75^th^ percentiles and median with whiskers representing minimum and maximum values.

## RESULTS

### Social Novelty Recognition is Impaired in SRKO mice

Patients with schizophrenia experience social withdrawal and impaired social and nonsocial recognition memory [34], and deficits in social cognition have been linked to altered E/I balance in the cortex [35, 36]. A three-chamber arena was used to assess sociability and social recognition by analyzing the proportion of time spent in each chamber and the number of nose-point entries within 5cm of each cage (**Figure 1 A**). During the sociability stage, EEG-tethered mice could investigate a novel mouse or a novel object. Behavioral measures of sociability were not significantly different between WT and SRKO mice. Both WT and SRKO mice spent a larger percent of time in the chamber containing the novel mouse than the chamber with the novel object [**Figure 1 B**, main effect of object, F(1,22)=7.543, p=0.0118], consistent with prior studies [25]. Using proportions (mouse/object) of the measurements described above, we directly compared the sociability preference between WT and SRKO animals. There were no significant differences between WT and SRKO animals in their preference for a novel mouse over a novel object (**Figure 1 D**, Welch’s t-test).

During the social novelty stage, mice could freely investigate the same mouse in the same location from the sociability stage (familiar mouse) or a novel mouse where the object had been in the prior stage. WT mice had a strong trend to spend a larger percent time in the novel mouse chamber than the familiar mouse chamber, while SRKO mice had a weak trend in the opposite direction [**Figure 1 E**, F(1,22)=8.171, p=0.0091, (WT) p=0.0576, (KO) p=0.0856]. We additionally measured the number of nose point entries in a 5cm zone surrounding each mouse or object as a complementary analysis. During the social novelty stage, only WT animals had more entries within 5cm of the novel mouse than the familiar mouse [**Figure 1 F**, F(1,22)=5.094, p=0.0343, p=0.0068]. Together, these results suggest WT but not SRKO animals spent more time investigating the novel mouse than the familiar mouse. The proportion of time spent in the novel/familiar mouse chamber was greater for WT (median=1.770) than for SRKO animals (median=0.505, Mann-Whitney U=6, p=0.0451, **Figure 1 G, left side**). Similarly, the number of nose point entries in a 5cm radius of the novel/familiar mouse was greater for WT (median =1.971) than for SRKO animals (median=0.7083, Mann-Whitney U=5, p=0.0295, **Figure 1 G, right side**). In summary, SRKO animals spent a smaller proportion of time investigating the novel versus familiar mouse compared to controls, indicating SRKO mice have decreased short-term social novelty recognition or impaired social novelty-related exploration. This is supported by another study using a three-chambered approach task in SRKO mice [37].

### Social Task-Elicited Gamma Power Is Impaired in SRKO mice

We additionally recorded social investigation associated gamma power from the frontal cortex during performance of the social novelty task. Because increases in gamma activity corresponded with the start of each novel mouse investigation (**Figure 2 A-B, D-E**), we focused our analysis on the first second of these investigations using 0.5s bins. During the sociability task, SRKO mice had a deficit in social task-elicited low gamma power (25-58Hz) compared to WT littermates from 0.5-1s after the novel mouse investigation began (**Figure 2 C**, t(372)=2.318, adjusted p=0.0415). Deficits in elicited gamma also emerged during the social recognition task. During the first second of novel mouse investigation, WT animals had a significantly larger increase in elicited low gamma power (25-58Hz) compared to SRKO animals (**Figure 2 F,** *0-0.5s*, t(332)=2.896, adjusted p=0.0040; *0.5-1s*, t(332)=3.490, adjusted p=0.0011). Elicited gamma was comparable between genotypes during the object and familiar mouse investigations (**Figure S1**).

**Figure 2.**
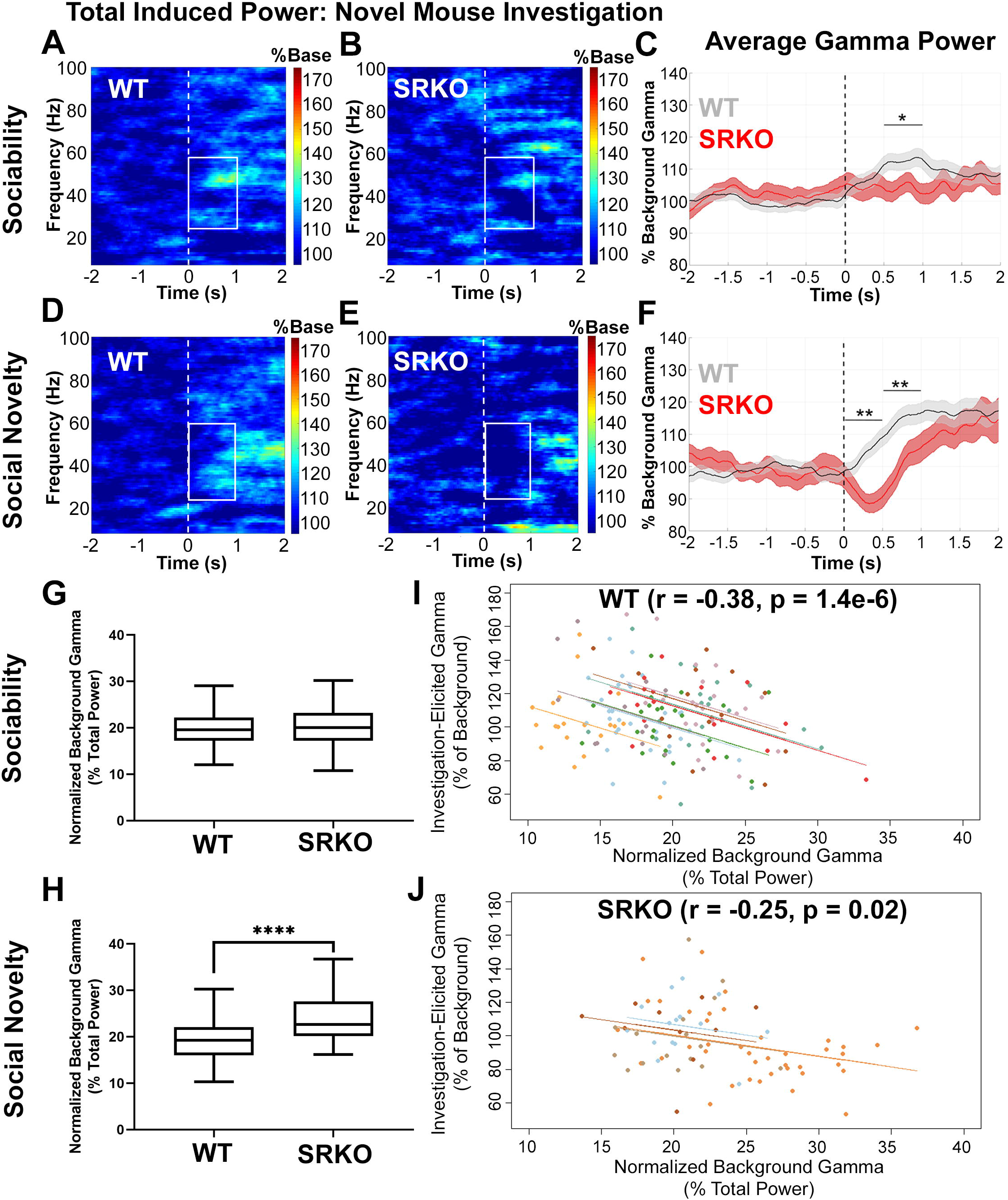
Social Task-Elicited Gamma Power Is Impaired in SRKO mice. Frontal cortex social task-elicited gamma power (25-58Hz) was recorded during the behavioral tasks described in Figure 1. Grand average spectrograms appear in **A-B** and **D-E** (% of baseline power), while the investigation-elicited gamma power normalized to background gamma is graphed in **C** and **F**. The dotted line represents the start of the novel mouse investigation (time 0) and the white boxes outline the data that was analyzed in 0.5s bins. WT mice had a significantly larger increase in gamma power than SRKO animals during 0.5-1s for the sociability task (**C**), and 0-1s for the social novelty task (**F**). Greater normalized background gamma power (0-4s prior to investigation, percent of total power) was evident in SRKO mice compared to WT littermates during the social novelty task (**H**), suggesting an improper signal-to-noise ratio may contribute to the difference in elicited gamma. Furthermore, there were significant inverse correlations such that greater background gamma power was associated with reduced investigation-elicited gamma power for all investigations across both trials for WT (**I**) and SRKO (**J**) animals. Each dot represents a single investigation of a novel object or a novel or familiar mouse, and each individual mouse is represented by a unique color. The data in Figure 1 and 2 suggest SRKO mice have a deficit in social recognition which corresponds with impaired social-elicited gamma in response to a novel mouse investigation. Enhanced background gamma in SRKO mice may be a contributing factor to their deficit in elicited gamma. Stars represent significance from multiple t-tests with Holm-Sidak correction (**C, F**) or a Mann-Whitney U test (**H**).

To determine if an improper signal-to-noise ratio of gamma power may contribute to this difference, we compared the normalized background broadband gamma power (0-4s pre-investigation, 25-58 Hz, percent of total power) between genotypes during novel mouse investigations. During the sociability stage, there was no difference in background gamma between genotypes (**Figure 2 G**).

However, during the social novelty stage, SRKO mice had significantly greater background gamma power (median=22.64) than WT littermates (median=19.24, **Figure 2 H**, Mann-Whitney U=1436, p<0.0001). Furthermore, there were significant inverse correlations between investigation-elicited gamma power and background gamma power for all investigations across both trials for WT (**Figure 2 I**; WT r= −0.38, p=1.4e-6) and SRKO animals (**Figure 2 J**; r= −0.25, p=0.02). Altogether, our data suggest SRKO mice have a deficit in social recognition, which corresponds with impaired social task-elicited gamma activity in response to investigating a novel mouse. Enhanced background gamma in SRKO mice may be a contributing factor to their deficit in elicited gamma, but this appears to be task specific. No changes in gamma power were observed during resting state activity in a separate context (**Figure S2, S3**).

### Sensory Gating is Impaired in SRKO mice

Sensory gating is an auditory processing phenomenon impaired in patients with schizophrenia and associated with hallucinations or delusions [7, 38]. The frontal cortex grand average ERPs generated by paired tones (S1 & S2) were examined (**Figure 3 A**). Although N40 was larger in S1 than S2 for all animals, only WT animals had significantly larger P20 and P20-N40 amplitudes in S1 than in S2 (p<0.05, **Figure 3 B,** <u>P20 interaction</u> F(1,17)=4.646, <u>N40 main effect</u> F(1,17)=18.33, **Figure 3 C,** <u>P20-N40 interaction</u> F(1,17)=5.213, see **Supplementary Materials B2**). Comparison of S2/S1 and S1-S2 “normalized” ratios revealed impaired sensory gating of P20-N40 amplitude in the frontal cortex of SRKO mice compared to WT littermates [**Figure 3 D**, <u>S2/S1</u>, t(13.39)=2.887, p=0.0124; **Figure 3 E**, <u>S1-S2</u>,t(15.40)=2.256, p=0.039].

**Figure 3.**
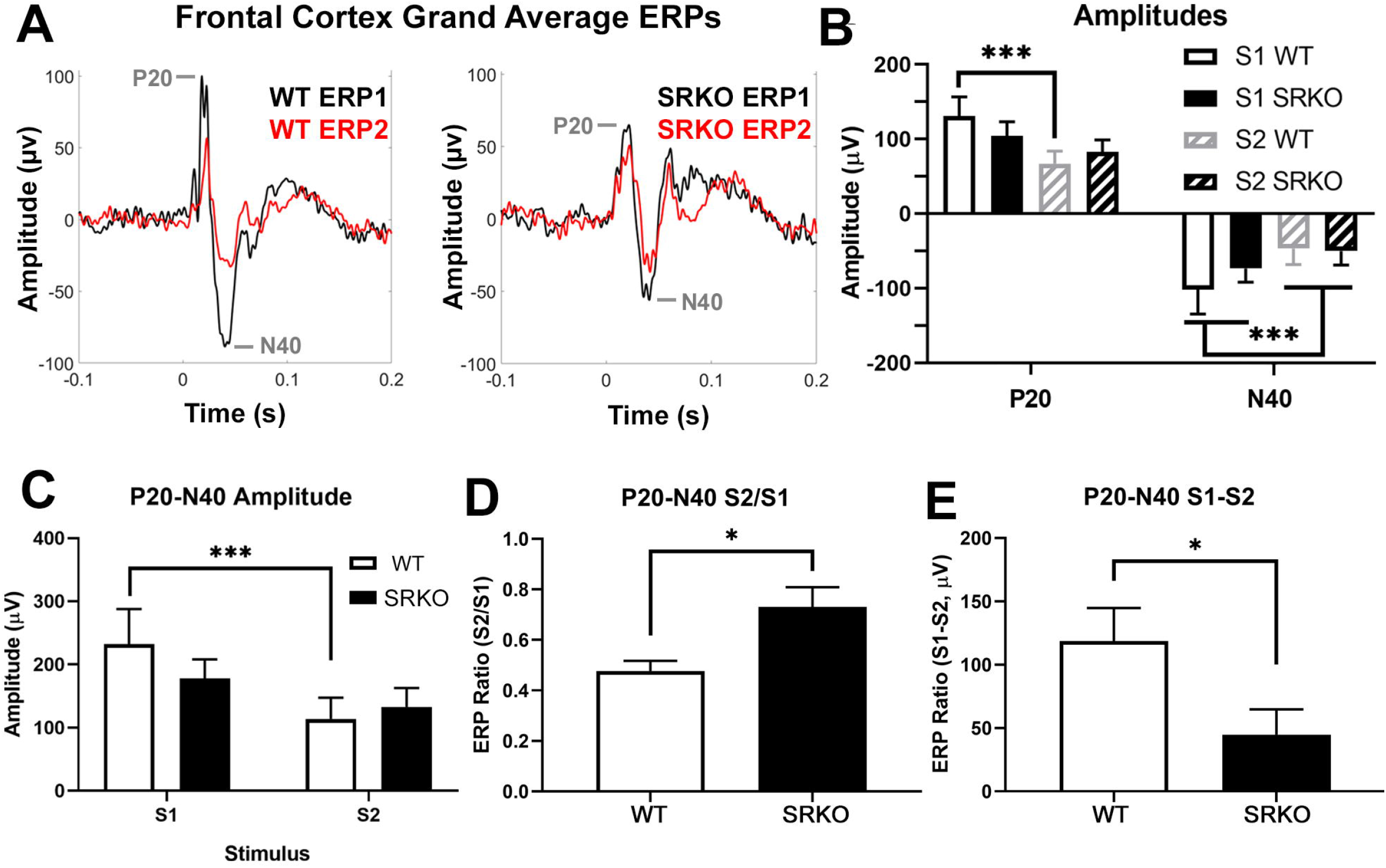
Sensory Gating is Impaired in SRKO mice. Sensory gating was measured in the frontal cortex EEG of SRKO mice (n=10) and WT littermates (n=9) by averaging the evoked response potential (**ERP**) of 100 repetitions of two identical 5kHz 50ms tones (S1 and S2) separated by 500ms. Frontal cortex grand average ERP’s are superimposed in **A** for WT (**left**) and SRKO (**right**) animals. Although N40 was larger in S1 than S2 for all animals, only WT animals had significantly larger P20 and P20-N40 amplitudes in S1 than in S2 (**B, C**). The P20-N40 amplitude was normalized as a ratio between ERP1 (from S1) and ERP2 (from S2) before comparing between groups. Compared to WT littermates, SRKO mice had a reduced gating response of P20-N40 amplitude in the frontal cortex as evidenced by a larger S2/S1 ratio (**D)** and an attenuated difference between S1 and S2 P20-N40 amplitudes (**E**). Stars in **B** and **C** represent a main effect of stimulus in a two-way RM ANOVA (N40) or significance in a Holm-Sidak post hoc test following a significant interaction in a two-way RM ANOVA (P20, P20-N40). Stars in **D** and **E** represent significance in an unpaired two-tailed Welch’s t-test.

Next, we examined whether there was a sensory gating deficit in gamma power (30-80Hz), as has been reported in patients with schizophrenia and a mouse model of schizophrenia [39]. As shown in **Figures 4 A and B,** KO mice also demonstrated impaired sensory gating of frontal cortex gamma power compared to WT [t(10.14)=2.634, p=0.0247]. Specifically, ERP2 gamma power was 75% reduced from ERP1 in WT mice, while it was only 42% reduced from ERP1 in SRKO mice. We additionally performed PSD comparisons between genotypes for S1 (ERP1) and S2 (ERP2) across the entire frequency range (0.5-100.5Hz; **Figure 4 C and D)**. Compared to WT, SRKO animals showed an overall decrease in power during ERP1 (**Figure 4 C**, 1-1.05s), but not ERP2 (**Figure 4 D**, 1.50-1.55s) [*<u>ERP1</u>* main effect of genotype, F(1,17)=4.576, p<0.0472].

**Figure 4.**
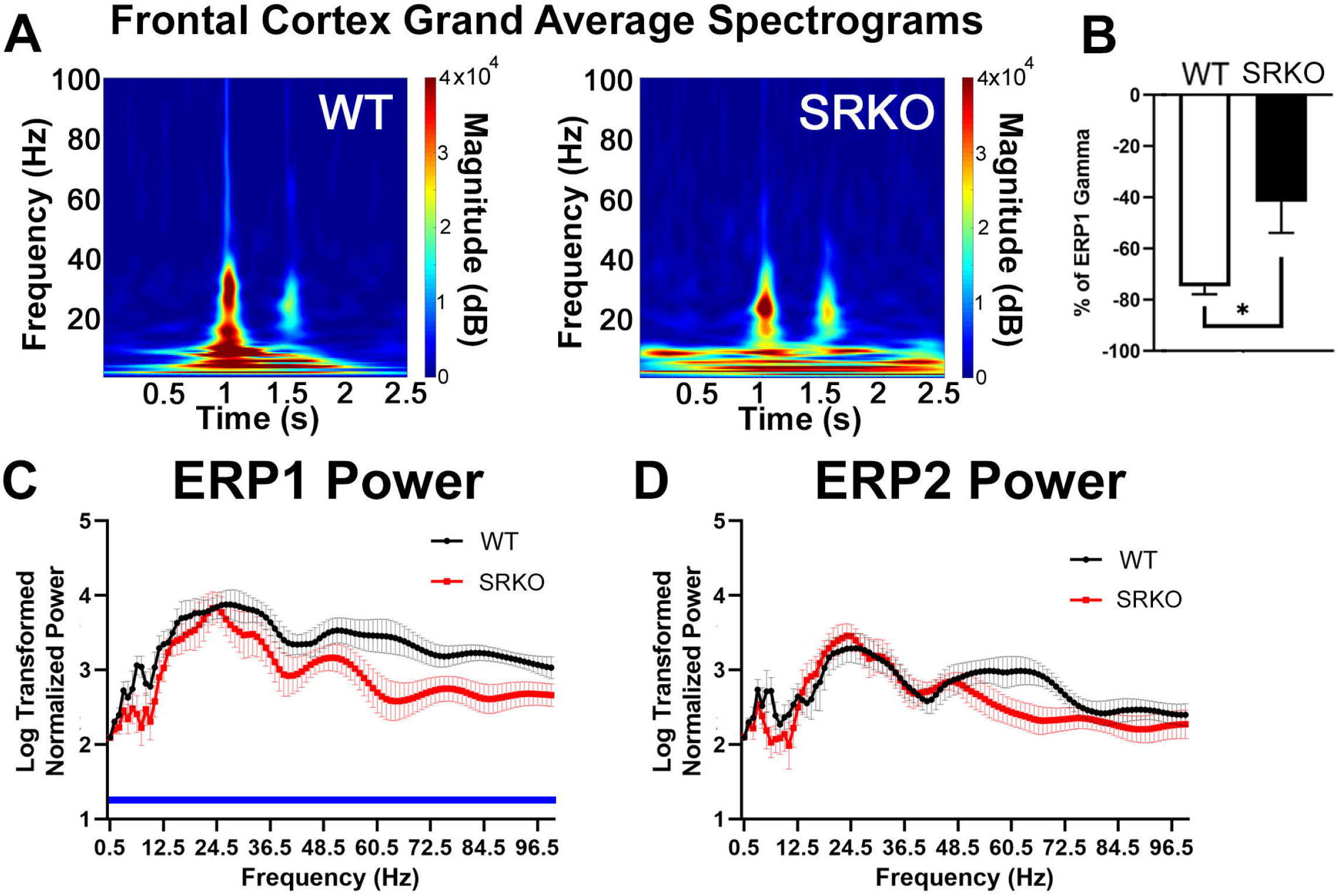
Power Spectral Density Abnormalities Occur in SRKO mice During Sensory Gating. We examined whether there was a sensory gating deficit in evoked gamma power (30-80Hz). The frontal cortex grand average spectrogram **(A)** demonstrates the change in evoked power during S1 (1-1.05s) and S2 (1.5-1.55s) for WT (**left**) and SRKO (**right**) mice. The frequency comparisons in parts **B-D** were calculated using these stimulus times. Similar to ERP amplitude, SRKO mice also demonstrated impaired sensory gating of gamma power (41.7% reduced from S1 to S2) compared to WT littermates (74.8% reduced, **B**). To examine task-evoked power spectral density differences between genotypes during each stimulus, the log-transformed normalized power for 0.5-100.5Hz was analyzed using 1Hz bins. Compared to WT littermates, SRKO animals had decreased power across the frequency spectrum during ERP1 (C) but not during ERP2 (D). Stars represent a significant unpaired two-tailed Welch’s t-test. C had a main effect of genotype in a two-way RM ANOVA, represented by a blue bar under the significantly different frequencies.

### Other Biomarkers are Neurotypical in SRKO mice

Other translationally-relevant electrophysiological biomarkers of schizophrenia investigated, including ASSR and sleep spindles, were unaltered in SRKO mice (see **Supplemental Materials A6, B3, and Table S1** for more detail). Elicited and induced (non-phase locked “background”) power and 40Hz phase locking were unchanged in the ASSR task between WT (n=11) and SRKO (n=10) mice (**Table S1**). There were no substantial changes between WT (n=14) and SRKO (n=12) animals for the percent time in each sleep/wake vigilance state, the average bout length, or average bout frequency. Sleep spindle characteristics (spindle density, amplitude, median and mean duration, median frequency) were not different between WT (n=7) and SRKO (n=6) animals. In the parietal and frontal cortices, significant differences in EEG frequency power spectra (<10 Hz) were found between SRKO mice and WT littermates during wake and sleep states in light and dark periods, although these effects were variable (see **Supplemental Figures S2 and S3**). Parietal cortex data was not significantly different between genotypes for most measurements (see **Supplemental Materials B4 and Figure S4**).

## DISCUSSION

SRKO mice have been well characterized to exhibit behavioral, brain morphological, and neurochemical abnormalities similar to what is observed in schizophrenia [22, 24, 25, 40–42].Here, we tested whether SRKO mice would phenocopy certain EEG biomarker abnormalities that are observed in patients with schizophrenia or induced by NMDAR antagonists. SRKO mice exhibited deficits in shortterm social recognition that corresponded with impaired investigation-elicited gamma power. Additionally, SRKO mice exhibited sensory gating impairments, in terms of both gamma power and ERP amplitude. However, other biomarkers such as ASSR and sleep spindles were unaffected in SRKO mice. Abnormal gamma power, whether task-associated or background, was a common theme across our study and may provide insight into the mechanisms behind these biomarker deficits.

### Deficient Task-Associated Gamma in SRKO Mice is Associated with Behavioral Impairments

E/I balance maintains stable yet flexible cortical activity, which manages a proper signal-to-noise ratio necessary for normal cortical function [36]. Deficits in GABAergic interneurons are common in many neuropsychiatric disorders and lead to abnormal gamma band oscillations at rest and during tasks [2]. This likely disturbs E/I balance through cortical disinhibition, leading to “noisier” circuits, and inefficient information processing [36]. Altered gamma oscillations in SRKO mice would suggest chronic NMDAR hypofunction disrupts the E/I balance with consequences on information processing. Indeed, SRKO mice had impaired task-associated gamma power in the frontal cortex during social recognition and sensory gating tasks which corresponded with deficits in task performance. Patients with schizophrenia often have impaired frontal cortex gamma band oscillations, which are associated with parvalbumin interneuron dysfunction and aberrant cognitive and perceptual functions [2]. Therefore, this deficit might be linked to previously reported reductions in cortical parvalbumin interneuron density in mature adult SRKO mice [27]. In addition, the impaired frontal cortex gamma power that occurred during novel social interactions, but not familiar mouse nor novel object investigations (**Figure S1**), could be a biomarker for social cognitive dysfunction that indicates a deficit in socially motivated working memory, attention, memory consolidation or retrieval.

We additionally observed that background (pre-investigation) broadband gamma power was abnormally high in the frontal cortex of SRKO mice during performance of the social novelty task (**Figure 2 H**), but not during independent measurement of gamma at rest (**Figure S2 A-B**). Interestingly, the default mode network (**DMN**) is a collection of brain regions including the prefrontal cortex with enhanced gamma activity during resting state behavior and suppressed gamma activity during cognitive tasks [43]. Patients with schizophrenia have an impaired ability to suppress the DMN, potentially because of an E/I imbalance, which contributes to working memory deficits and other cognitive impairments [44]. Thus, our findings suggest that SRKO mice may have deficient suppression of background gamma activity during certain behavioral contexts (social investigation) that could act as excess “noise” that disrupts the signal-to-noise ratio and cortical E/I balance, which contributes to impaired task performance.

### Sensory Gating Deficits in SRKO Mice are Similar to Schizophrenia

Our sensory gating findings revealed multiple similarities between SRKO mice and patients with schizophrenia. Patients with first episode or chronic schizophrenia have a lower S1 P50 amplitude, a comparable or larger S2 P50 amplitude, a larger S2/S1 ratio, and a smaller S1-S2 difference during sensory gating [5, 45–47]. In our study, WT animals had significant gating of frontal P20 and P20-N40 amplitude (the mouse analogues of the human P50) [48, 49]. Therefore, SRKO mice had a larger S2/S1 ratio and a smaller S1-S2 difference similar to what is often seen in patients with schizophrenia. Furthermore, SRKO mice had deficits in frontal cortex evoked power across the gamma frequency range during sensory gating consistent with the clinical schizophrenia literature and another a genetic mouse model relevant to schizophrenia [39]. Normalized beta and gamma power were among frequencies lower in SRKO mice than WT littermates during the period when the ERP1 P20 and N40 occur (0-50ms). Comparable deficits in beta (20-30Hz) and gamma power (30-50Hz) were reported in patients with schizophrenia for the human analogues of these peaks (ERP1 P50 and N100, 0-100ms), and may be a biomarker of a failed initial sensory registration [45].

Although impaired sensory gating is observed in patients with schizophrenia, pharmacologically-induced NMDAR hypofunction does not usually affect this biomarker. Neither acute nor chronic administration of the NMDAR antagonists ketamine or MK-801 significantly impairs sensory gating (S2/S1 ratio) in humans, mice, or rats [50–54]. Furthermore, mice with reduced NMDAR NR1 subunit expression have a normal sensory gating response despite increased P20 and N40 amplitudes [55], but another group reported a reduced S2/S1 ratio in these mice [56]. Altogether, this suggests the sensory gating deficits observed in SRKO mice may not be due to NMDAR hypofunction alone. Low cortical dopamine levels may contribute to a sensory gating deficit since ketamine (which induces dopamine release [57]) does not induce these deficits [52] and most antipsychotics with D2-antagonist properties fail to rescue these deficits [58, 59]. Mice with reduced alpha-7 nicotinic acetylcholine receptors have sensory gating impairments [39], and clozapine (which enhances acetylcholine [60]) and nicotine each rescue sensory gating deficits in patients with schizophrenia or their relatives [58, 59, 61, 62]. Therefore, reduced cholinergic or cortical dopaminergic activity, as seen in schizophrenia [63, 64], may contribute to the sensory gating deficits in schizophrenia and SRKO mice.

### ASSR and Spindles are unaffected in SRKO mice

Synchronization of cortical neuron firing in response to repetitive stimuli is believed to depend critically on E/I balance and represents a process related to cognitive function [65]. Patients with schizophrenia and rodents administered NMDAR antagonists exhibit abnormalities in their ability to synchronize cortical firing in response to 40Hz auditory stimuli [32, 66–68]. However, consistent with recently published findings [69], ASSR evoked power was intact in SRKO mice at all measured frequencies (**Table S1**). Phase locking and background (induced) power were also unchanged at 40Hz (**Table S1**). Acute versus chronic NMDAR antagonist treatments have competing effects that may explain our results. In tethered, freely moving rats acute MK-801 altered the intertrial coherence of the 40Hz ASSR in the primary auditory cortex, but chronic (21 day) MK-801 treatment had no significant effects [70]. Furthermore, the consequences of ketamine on the 40Hz ASSR in conscious rats depends largely on the dose used, the degree of NMDAR occupancy, and the amount of time since drug administration [71]. Therefore, our neurotypical ASSR result in SRKO mice is compatible with chronic NMDAR antagonist pharmacological studies. This contrasts deficient task-evoked gamma power in other biomarkers; however, these results may arise from distinct mechanisms. Indeed, the 40Hz ASSR deficit in patients with schizophrenia may be due to enhanced background gamma activity [32], which our mice do not replicate during this task or at rest (**Figures S2, S3**).

Region specificity could explain why biomarker deficits were seemingly inconsistent and mainly limited to the frontal cortex. The relative expression of SR, D-serine, and glycine in various brain regions could influence NMDAR signaling and the existence or location of biomarker deficits in SRKO mice. If D-serine and glycine coexist in a brain region, the level of activity can influence which one is used as an NMDAR co-agonist [72]. SRKO mice had no deficits in the 40Hz ASSR or sleep spindles, which are biomarkers that require intact thalamocortical circuitry [73, 74] or normal thalamic NMDAR signaling [75, 76]. Regions like the thalamic reticular nucleus do not express high levels of SR ([77, 78]; **unpublished data**) and thus likely have normal NMDAR function in SRKO mice. Furthermore, NMDAR antagonists have region-specific consequences on gamma power. MK-801 enhances gamma activity in the hippocampus and above bregma but not in the primary auditory cortex of conscious, freely-behaving rats [54]. Conversely, bath application of ketamine onto rat coronal brain slices enhances gamma rhythms (30-50Hz) in the primary auditory cortex, impairs gamma rhythms in the medial entorhinal cortex, and has no effect on hippocampal slices [79]. Therefore, gamma power in our parietal and frontal electrodes should not necessarily align in our global mouse model of NMDAR hypofunction. Future studies deleting SR in discrete cortical brain regions will address this confound.

### NMDAR Hypofunction Changes Delta and Theta Power

Several observed changes in PSD aligned with NMDAR hypofunction. Chronic NMDAR antagonist treatment can reduce theta and gamma power for weeks or months after the last drug administration in rodents [80, 81]. This may explain the SRKO deficit in broadband power during sensory gating and the reduction in parietal cortex normalized power around the theta band (4-7Hz) during resting state and sleep behaviors for the lights-off (active) period (**Figure S3**). We also found small differences in EEG delta power (1-4 Hz) during spontaneous NREM sleep, a frequency band that has been reported to be dysregulated with schizophrenia and is involved in cognition [82, 83], which suggests that NMDAR hypofunction could be involved in certain cognitive impairments observed with schizophrenia.

### Limitations

All animals expressed a left side preference (% time in chamber) during the empty three chamber habituation task, but the number of chamber entries were equal for left and right sides (See **Supplemental Materials B5**). The influence of baseline side preference on sociability and social novelty should equalize as the side of stimulus presentation was counterbalanced.

### Conclusion

In conclusion, the SRKO mouse model mimics a subset of EEG and behavioral phenotypes associated with schizophrenia and chronic NMDAR antagonist treatment. These novel biomarker deficits compliment SRKO literature reporting changes similar to positive, negative, and cognitive symptoms of schizophrenia [22–26, 69]. These mice may be useful for modeling patients with chronic schizophrenia more accurately than pharmacologic models in specific domains [84] including sensory gating. Future studies of SRKO mice can confirm whether 1) these phenotypes persist among antipsychotic or D-serine treatment, 2) glycine-related compensatory responses are occurring, 3) thalamocortical circuitry is intact, or 4) abnormalities in dopamine levels, cholinergic signaling, or parvalbumin-containing neurons exist in the neocortex. Our gamma band findings support the idea of an E/I imbalance manifested as an aberrant signal-to-noise ratio impairing cognition and information processing. This deficit may be tied to impaired task-dependent alteration in functional connectivity and impaired suppression of the DMN. Understanding the mechanisms behind these biomarkers could lead to personalized early interventions that prevent the transition to psychosis.

## Acknowledgements

We would like to thank Dr. Robert W. McCarley for his efforts in forming this productive collaboration, and Yunren Bolortuya for assistance with EEG implantation and sleep scoring.

## Author Contributions

D.D.A., and J.M.M. conceived and designed the experiments; D.D.A., L.K.R., F.L.S. and J.M.M. performed the experiments and analyzed the data; D.D.A and J.M.M. drafted and revised the manuscript for content. Others: conceived the experiments, interpreted results, and revised the manuscript.

## Supplemental Material

accompanies this paper at Biorxiv.org. Supplemental citations include [85, 86].

