## Supplemental Material for "Altered neural oscillations and behavior in a genetic mouse model of NMDA receptor hypofunction"

The Supplemental Material includes Supplemental Methods (**A**), Supplemental Results (**B**), Table S1, and Figures S1-S4.

**A. Supplementary Methods.**

*A.1. Automated Video Tracking:* Video tracking was set to trigger a TTL event (marking a putative investigation) if the animal's nose point reached within 5cm of a specific cage, and ended when the animal's nose was over 10cm away from the cage. The two cages had distinct TTL events to distinguish between animal/object or familiar/novel mouse investigations.

*A.2. Sensory Gating Data Analysis:* Raw EEG records were averaged over 100 paired-tone presentations to create a waveform average for each animal that contained one average trial (two stimuli). An epoch of -100 to 400ms around each stimulus was extracted from this average and baseline corrected to the mean of their 100ms before stimulus onset. For all baseline corrections (sensory gating and MMN), we subtracted all amplitude values within an epoch from the mean amplitude value of a pre-stimulus baseline (in this case 0 to 100ms before stimulus onset). For each ERP we measured the maximum positive deflection around 20ms (P20, 15-30ms after the tone) and the maximum negative deflection around 40ms (N40, 25-55ms).

A continuous wavelet transformation was applied to the waveform average to calculate evoked spectral power from 0.5 to 100.5Hz in 0.5Hz steps across an epoch of 3 seconds which encompassed both stimuli. Averaged gamma (30-80Hz) power was normalized to a 0-500ms pre-stimulus "baseline" period and summed over the periods when each stimulus was audible.

Averaged gamma (30-80Hz) power was normalized by dividing each time point (representing 0.0005 seconds) by the average of the gamma power during a 0-500ms pre-stimulus "baseline" period. This normalized power was then summed across the time each ERP was audible (ERP1: 1-1.05 seconds; ERP2 1.5-1.55 seconds) and the percent change in gamma power from ERP1 to ERP2 was calculated. The complete frequency band (0.5-100.5Hz) was also analyzed by averaging into 1Hz frequency bins. This data was normalized and summed in the same way described above, only comparing within identical frequency bins.

*A.3. Sensory Gating Threshold:* Mice were removed from analysis if they had no detectable ERP, using the following criteria: ERP1 P20 amplitude was less than 1.25 times the peak during a baseline period 50-100ms before the tone, or if ERP1 or ERP2 had an N40 that occurred before P20. In the gamma analysis, one WT mouse had its parietal EEG data removed for being a significant outlier among the data (Grubb's test for outliers, $Z=2.63$, $p<0.05$).

*A.4. ASSR:* This task measures cortical entrainment to repetitive auditory stimuli. Following a 15 minute habituation, a 1s train of audible clicks (90dB) is presented at a range of frequencies in a counterbalanced pseudorandom order (20-60Hz in 10Hz steps, 300 reps per frequency) while cortical EEG response is measured. Evoked power is the average power (stimulus frequency ±2Hz) during the stimulus (1.25-2s) divided by the average power during a baseline period (0-0.75s). Induced power is measured identically but uses non-phase locked power. Phase locking value is a measure of phase synchronization for each stimulation frequency during the period of stimulation. To calculate 40Hz phase locking value, a wavelet transformation was applied to the 40Hz stimulation data and the phase angle of 37-43Hz was calculated for each trial. For all ASSR experiments, there were 11 WT mice (8 males, 3 females) and 10 KO mice (7 males, 3 females).

*A.5. Statistical Analyses:* The effect of subjects ("matching") in the two-way RM ANOVA was significant unless mentioned. For more detail on spectral analysis, see the sensory gating section above.

*A.6. Sleep / Wake Data*: Freely moving EEG-tethered male and female SR KO mice (n=12) and WT littermates (n=14) habituated for 48h and were recorded for 24h. Researchers blinded to genotype manually sleep scored EEG data in 4s epochs using Sirenia software (Pinnacle Technology). A change in the reference electrode location from the contralateral parietal cortex to the cerebellar cortex occurred halfway through the experiment (n=7 WT 6 KO mice per reference electrode location). This had the potential to affect power spectral density and sleep spindle measurements, so only cerebellar referenced animals were included in these analyses. Sleep spindles were detected using an automated MATLAB-based algorithm developed in-house (Uygun et al. 2019 *SLEEP*). Custom MATLAB scripts were used to examine power spectral density. A multitaper spectral analysis (Chronux toolbox, http://chronux.org/; 2 tapers, bandwidth 3) was performed on each 4s scored epoch (Mitra and Bokil 2007). Within each behavioral state (wake, NREM, REM), each epoch was normalized to the sum power of all epochs (0–100 Hz) in the baseline day hours 0–24. See **Supplementary Figures S2 and S3** for power spectral density results, and **Supplementary Materials B3** for Sleep/Wake experimental results.

**B. Supplementary Results**

*B.1. Means ± Standard Error of the Mean:* The values of individual treatment groups are presented here for experiments that had significant or near-significant results.

| Figure | WT Value | SRKO Value |
|---|---|---|
| **1B** | Mouse 39.03 ± 3.18%,<br>Object 33.89 ± 4.49% | Mouse 45.07 ± 4.13%,<br>Object 26.58 ± 4.85% |
| **1E** | Novel 46.72 ± 5.80%,<br>Fam 28.31 ± 5.05% | Novel 25.20 ± 8.79%,<br>Fam 43.17 ± 5.60% |
| **1F** | Novel 23.50 ± 2.11 entries,<br>Fam 10.63 ± 1.76 entries | *KO values were not significant (n.s.)* |
| **2C** | **0-0.5s:** n.s.<br>**0.5-1s:** 112.6 ± 2.4% | **0-0.5s:** n.s.<br>**0.5-1s:** 104.1 ± 2.7% |
| **2F** | **0-0.5s:** 103.3±2.2%<br>**0.5-1s:** 114.1±2.7% | **0-0.5s:** 91.2±2.6%<br>**0.5-1s:** 99.5±3.4% |
| **2H** | 19.20 ± 0.39%;<br>Median value 19.24% | 24.05 ± 0.68%;<br>Median value 22.64% |
| **3B P20 (Left)** | **S1**: 130.8 ± 25.66 µv<br>**S2**: 67.16 ± 16.60 µv | **S1**: 104.2 ± 18.90 µv<br>**S2**: 82.89 ± 15.99 µv |
| **3B N40 (Right)** | **S1**: -101.8 ± 32.85 µv<br>**S2**: -46.68 ± 21.38 µv | **S1**: -73.48 ± 18.25 µv<br>**S2**: -50.08 ± 18.71 µv |
| **3C** | **S1**: 232.6 ± 55.31 µv<br>**S2**: 113.8 ± 33.72 µv | **S1**: 117.7 ± 30.53 µv<br>**S2**: 133.0 ± 29.77 µv |
| **3D** | 0.4772 ± 0.04034% | 0.7305 ± 0.07793% |
| **3E** | 118.8 ± 26.06% | 44.76 ± 19.91% |
| **4B** | -74.8±3.1% | -41.7±12.2% |
| **4C (0-100Hz average)** | 3.621 ± 0.042 | 3.392 ± 0.055 |

*B.2. Additional Statistics and Comparisons for Figure 3 Sensory Gating*. Sensory gating deficits occurred in the frontal cortex of SRKO mice (n=10) in comparison to WT littermates (n=9). For all animals, peak amplitudes of the evoked response potential over the frontal cortex were larger during stimulus 1 (S1) than stimulus 2 (S2) during the sensory gating task [two-way RM ANOVA, main effect of stimulus; *P20* **(Fig 3 B)**, F(1,17)=18.78, p=0.0005; *N40* **(Fig 3 B)**, F(1,17)=18.33,p=0.0005; *P20-N40* **(Fig 3 C)**, F(1,17)=25.45, p<0.0001]. However, there was also an interaction such that only WT mice had a larger P20 **(Fig 3 B)** and P20-N40 **(Fig 3 C)** in S1 compared to S2 [two-way RM ANOVA, *P20 interaction* F(1,17)=4.646, p=0.0457, Holm-Sidak p=0.0007 for WT S1 vs WT S2; *P20-N40 interaction* F(1,17)=5.213, p=0.0356, Holm-Sidak p=0.0002 for WT S1 vs WT S2].

ERP ratios were also calculated, and there was a larger P20-N40 S2/S1 ratio **(Fig 3 D)** and a smaller P20-N40 S1-S2 ratio **(Fig 3 E)** for KO animals [unpaired two-tailed Welch's t-test, P20-N40 S2/S1 t(13.39)=2.887, p=0.0124; P20-N40 S1-S2 t(15.40)=2.256, p=0.039].

Similar to the ratios of P20-N40 amplitude, ratios of P20 amplitude revealed impaired sensory gating of P20 amplitude in the frontal cortex of SRKO mice compared to WT littermates [*data not shown,* P20 S2/S1, t(17)=2.813, p=0.015; WT 0.522±0.057 arbitrary units, SRKO 0.905±0.124 a.u.; P20 S1-S2, t(17)=2.169, p=0.045; WT 63.64±13.39 µv, SRKO 21.36±14.17 µv].

There were no differences in the N40 S1-S2 and N40 S2/S1 ratios (*data not shown*).

*<u>B.3. Sleep and Spindle Results:</u>* There were no substantial changes between genotypes in the following measurements:

- Percent time in each vigilance state (wake / NREM / REM, 1hr bins), n=14 WT and 12 KO
- Average bout length and average bout frequency for each vigilance state (wake / NREM / REM, 1hr bins), n=14 WT and 12 KO
- NREM sleep spindle density, amplitude, median and mean duration, median frequency (1hr bins, n=7 WT and 6 KO animals
    - Mean spindle duration was elevated in SRKO mice for a few hours after the transition to the lights-off period *(data not shown)*

*<u>B.4. Parietal Cortex Supplementary Results:</u>* There were no significant changes between genotypes for the following measurements:

- Social task-evoked gamma power
- Sensory gating peak amplitudes and ratios (see **Supplementary Figure S4**)
- Sensory gating change in evoked gamma power or power spectral density
- ASSR evoked power (20-60Hz) or induced power (40Hz)
- NREM sleep spindle characteristics

*<u>B.5. Side Preference in three chamber habituation.</u>* During the empty three chamber habituation before the sociability task, all animals expressed a side preference for the left chamber (two-way RM ANOVA, variables were genotype and location, main effect of location, $F(1,11)=8.432$, $p=0.0143$, Left side 39.77 ± 2.41% of total time, Right side 26.42 ± 2.92% of total time). However, the number of chamber entries during the habituation were not significantly different for left and right sides (two-way RM ANOVA, $p>0.05$). The influence of baseline side preference on sociability and social novelty should equalize as the side of stimulus presentation was counterbalanced.

**Supplementary Table S1. Frontal Cortex ASSR Values.**

| Stimulation Frequency | Evoked | Induced | Phase Locking |
|---|---|---|---|
| **20 Hz** | WT 45.79 ± 12.83;<br>KO 36.79 ± 6.489 | WT 0.9478 ± 0.0353;<br>KO 1.001 ± 0.0252 | |
| **30 Hz** | WT 48.94 ± 13.09;<br>KO 43.79 ± 12.67 | WT 0.9551 ± 0.0199;<br>KO 0.9581 ± 0.0220 | |
| **40 Hz** | WT 41.27 ± 8.212;<br>KO 51.06 ± 11.09 | WT 1.009 ± 0.0252;<br>KO 1.022 ± 0.0145 | WT 0.2874 ± 0.0374;<br>KO 0.3036 ± 0.0304 |
| **50 Hz** | WT 45.72 ± 22.09;<br>KO 26.25 ± 3.201 | WT 1.031 ± 0.0228;<br>KO 1.032 ± 0.0158 | |
| **60 Hz** | WT 43.03 ± 17.93;<br>KO 15.27 ± 3.769 | WT 1.053 ± 0.0136;<br>KO 1.016 ± 0.0163 | |
| **40Hz Harmonic (20Hz stim)** | WT 20.19 ± 5.439;<br>KO 25.89 ± 8.469 | | |

Average values per genotype (stimulation / background) are reported in this table. During the ASSR task, elicited power around each stimulus frequency (+/- 2Hz) was not different for any stimulus frequency between WT (n=11) and KO mice (n=10, two-way repeated measures ANOVA). The induced (non-phase locked) power was also unchanged between WT and KO mice during all stimulation frequencies (two-tailed unpaired Welch's t-test). Finally, 40Hz phase locking was compared between WT and KO animals but again there were no significant differences between groups (two-tailed unpaired Welch's t-test).

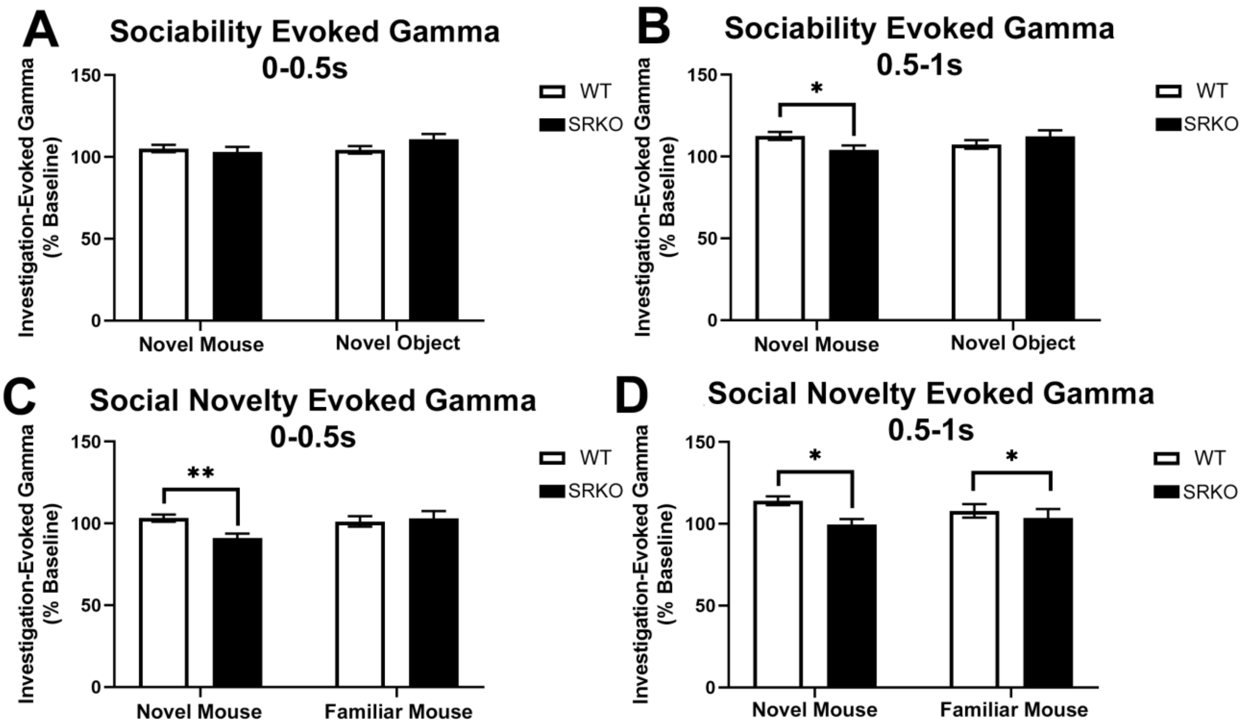


**Supplementary Figure S1. Object and familiar mouse investigations did not generally evoke differences in investigation-evoked gamma.** As reported in Figure 2, novel mouse investigations generate higher evoked gamma power in WT (n=8) than KO mice (n=5) during both the sociability (**B,** 0.5-1s of investigation) and social novelty stages (**C-D,** 0-1s of investigation). Evoked gamma power is similar between genotypes for object **(A, B)** and familiar mouse investigations **(C)**. Although there was a difference in gamma power during familiar mouse investigations in **D**, this was likely driven by novel mouse investigations (two-way ANOVA, main effect of genotype). Data is presented as mean +/- standard error of the mean. Stars represent: **B, C:** two-way ANOVA, significant interaction, Holm-Sidak $p<0.05$**, D:** two-way ANOVA, main effect of genotype $p<0.05$

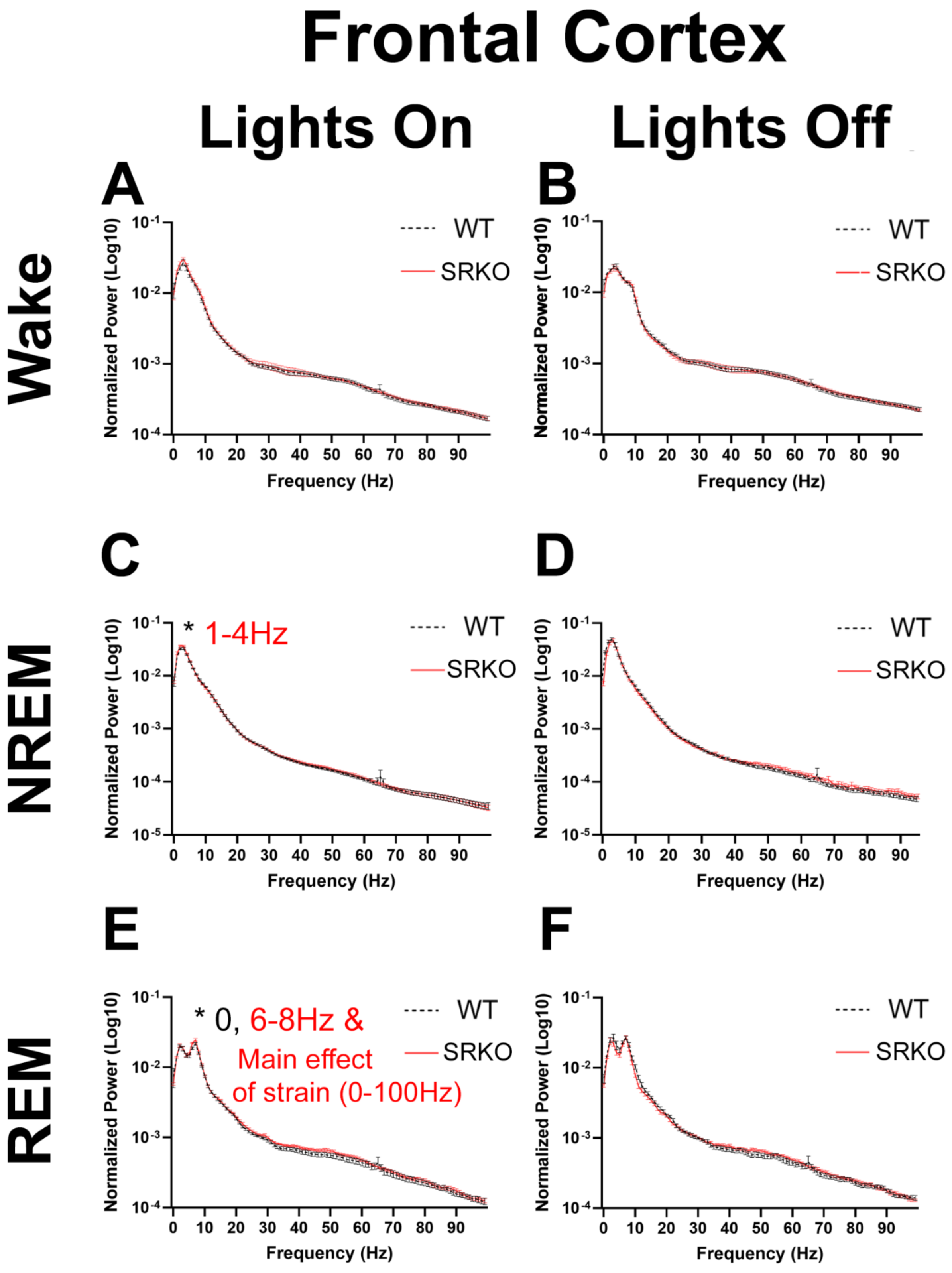


**Supplementary Figure S2. No change in resting state gamma power in the frontal cortex of SR KO mice.** Conscious, freely moving mice were housed at room temperature and given food and water ad libitum on a 12hr light / dark schedule (lights-on 7am, lights-off 7pm). Following 48hrs of tethered habituation, 7 WT and 6 SR KO mice were recorded for 24hrs. The normalized power spectral density during wake, non-rapid eye movement (NREM) and rapid eye movement (REM) sleep were analyzed separately in 1Hz frequency bins from 0-100Hz. Within each behavioral state (Wake / NREM / REM), each 4-second wake epoch was normalized to the sum power of all epochs (0-100Hz) across the 24hr recording.

There were no changes during wake, but during the lights-on (inactive) period some differences arose during NREM and REM sleep for frequencies under 10Hz. The starred text on each graph signifies frequency ranges that were significantly different between groups (two-way RM ANOVA, genotype x frequency interaction $p<0.05$, Holm-Sidak post hoc $p<0.05$). The color of the starred text (black for WT, red for KO) signifies the group with higher power at that frequency range. During REM sleep, KO animals also had higher power than WT littermates across the entire frequency band during the lights-on period (two-way RM ANOVA, main effect of genotype).

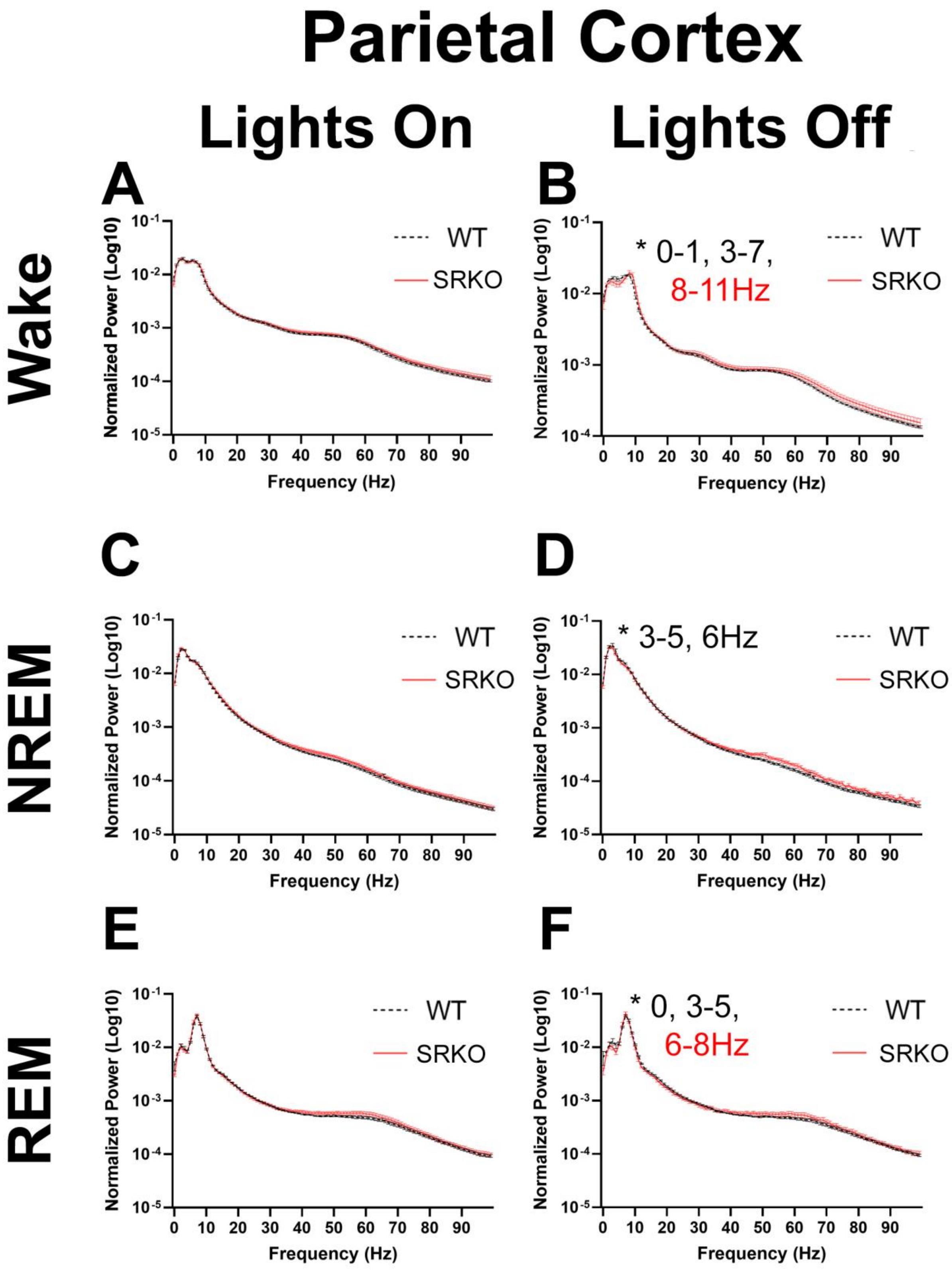
Parietal Cortex
Lights On
Lights Off
Wake
NREM
REM
A
B
C
D
E
F
WT
SRKO
* 0-1, 3-7,
8-11Hz
* 3-5, 6Hz
* 0, 3-5,
6-8Hz
Normalized Power (Log10)
Frequency (Hz)

**Supplementary Figure S3. Reduced delta and theta power in the parietal cortex of SR KO mice.** See **Figure S2** and **Supplementary Materials A6** for experimental details. The normalized power spectral density during wake, NREM and REM sleep were analyzed separately in 1Hz frequency bins from 0-100Hz. During the lights-off (active) period, normalized power around the delta (0.5-4Hz) and theta (~4-7Hz) bands were lower in SRKO mice than WT littermates during wake, NREM, and REM sleep. The starred text signifies frequency ranges that were significantly different between groups (two-way RM ANOVA, genotype x frequency interaction $p<0.05$, Holm-Sidak post hoc $p<0.05$). The color of the starred text (black for WT, red for KO) signifies the group with higher power at that frequency range.

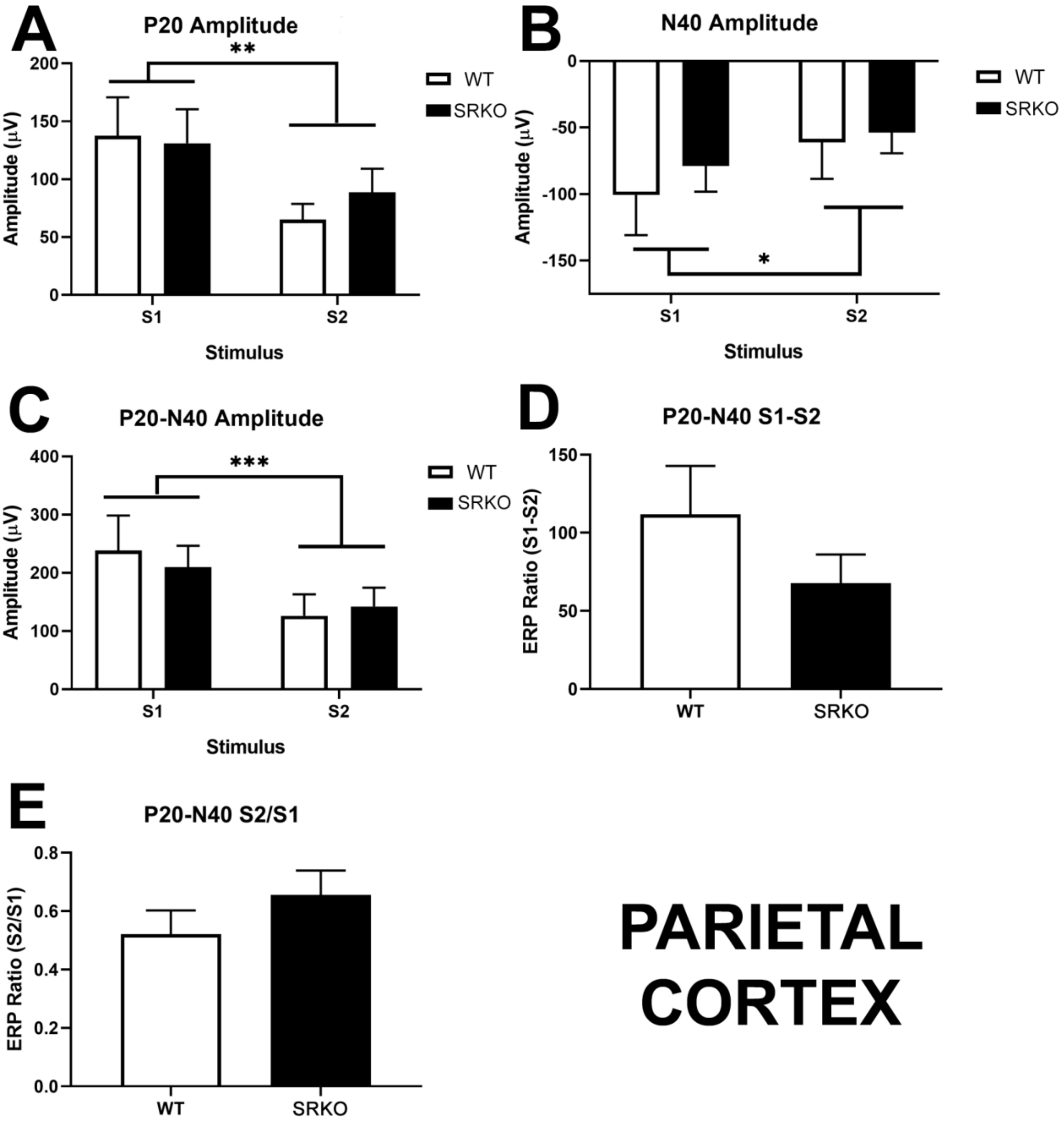


**Supplementary Figure S4. Sensory gating is neurotypical in the parietal cortex.** For all animals, peak amplitudes of the evoked response potential over the parietal cortex were larger during stimulus 1 (S1) than stimulus 2 (S2) during the sensory gating task [two-way RM ANOVA, main effect of stimulus; *P20* **(A)***,* F(1,16)=11.22, p=0.0041; *N40* **(B)***,* F(1,16)=8.189, p=0.0113; *P20-N40* **(C)***,* F(1,16)=24.88, p=0.0001]. No WT vs KO differences were observed between the following ratios: P20-N40 S1-S2 **(D)**, P20-N40 S2/S1 ratio **(E)**, N40 S1-S2 (*not shown*), N40 S2/S1 (*not shown*). Sample size was 9 WT 9 KO animals in the parietal cortex.
